# Mitochondrial-Targeted HDAP2 Preserves Retinal Ganglion Cells and Increases Pressure Tolerance in DBA/2J Mice

**DOI:** 10.1101/2025.10.23.684211

**Authors:** Margaret A. MacNeil, Widnie Mentor, Alexander Birk

## Abstract

**Purpose:** Retinal ganglion cell (RGC) loss in glaucoma occurs in a large fraction of patients even after intraocular pressure (IOP) is reduced. Mitochondrial dysfunction is a key mechanism that links elevated IOP to RGC degeneration. We tested whether HDAP2, a novel high-density aromatic peptide that binds cardiolipin to stabilize mitochondrial membranes, can protect RGCs in the DBA/2J mouse model.

**Methods:** DBA/2J mice received HDAP2 (3 mg/kg, intraperitoneally, every other day) starting at 4 months of age for 8 months. IOP was measured each month to track pressure exposure. RGC survival was assessed by counting RBPMS-stained cells in retinal wholemounts and optic nerve axons in semithin toludine blue-stained sections.

**Results:** HDAP2-treated retinas had ∼49% more RGCs than untreated retinas at similar pressure exposures (p = 0.0063; F(2,59) = 5.524). At mild IOP exposure, HDAP2 preserved 58% more RGCs compared with untreated retinas, and at high IOPs, RGC survival was 180% greater.

Kaplan–Meier analysis indicated that HDAP2 increased the threshold for severe RGC loss by 29 mmHg and reduced the chance of developing severe RGC degeneration by a factor of 4.6. Optic nerve axons from treated retinas were also well preserved, with axon morphologies appearing indistinguishable from controls. Axon size distributions did not change significantly among the treatment groups, suggesting that protection by HDAP2 was similar among RGC subtypes.

**Conclusions:** HDAP2 preserved both RGCs and axons in a pressure-dependent manner and increased tolerance to IOP. These results suggest that HDAP2 may complement pressure-lowering therapy, including for normal-tension and treatment-refractory glaucoma.

## Introduction

Glaucoma is a leading cause of irreversible blindness from the progressive degeneration of retinal ganglion cells (RGCs) and their axons. Increased intraocular pressure (IOP) is the primary risk factor ^1^, but many patients continue to lose vision despite reduced IOP ^2,3^. This fact highlights the necessity of developing therapies that enhance RGC tolerance to elevated intraocular pressure.

Mitochondrial dysfunction is a key mechanism that links elevated IOP to the death of RGCs observed in glaucoma ^4,5^. RGCs are uniquely vulnerable to metabolic distress that is driven by increased IOP. The cells’ unique architecture, with large somas, extensive dendritic arborizations, dense synaptic inputs, and unmyelinated intraretinal axons, requires high concentrations of mitochondria, typically found in compartments with the highest energy needs ^6–8^. Elevated IOP exerts mechanical stress on unmyelinated RGC fibers when leaving the eye, which disrupts transport of mitochondria and trophic factors. At the same time, compression of retinal blood vessels reduces oxygen availability, which leads to ischemia, increased reactive oxygen species (ROS) production, and oxidative damage ^9^.

When mitochondria generate ATP to support neuronal activity, the process of oxidative phosphorylation naturally produces ROS, which at physiological levels, is used for intracellular signaling. However, when ATP output is accelerated or a cell undergoes ischemic stress, excess ROS is generated ^10^, which can lead to damaged mitochondrial DNA, induction of the mitochondrial permeability pore (MPTP), disrupted mitochondrial quality control mechanisms, and reduced ATP output ^11–13^. In glaucoma, excess ROS accumulation has been correlated with mitochondrial fission and cristae depletion ^14^, caspase activation, and ultimately RGC apoptosis^5,9,15^. Thus, even minor perturbations in mitochondrial function can greatly influence RGC survival.

Mitochondrial dysfunction emerges early in glaucoma and plays a critical role in RGC degeneration. Several studies have shown that disruptions in axonal transport, reduced energy production, and structural changes in mitochondria precede measurable cell death ^16–18^. Similar patterns emerge in ischemia models, where mitochondrial failure and ATP depletion accelerate injury ^19^. Age further exacerbates this vulnerability, as mitochondrial decline in DBA/2J mice heightens RGC sensitivity to elevated IOP. Nicotinamide supplementation provided strong protection against these deficits and preserved RGC survival, although continuous treatment was necessary to maintain the benefit ^20^. Other mitochondria-directed approached have been tested, including antioxidants such as SS-31 and MitoQ, which reduce oxidative stress ^21,22^, as well as experimental approaches such as mitochondrial transplantation ^23^. While these interventions provide some benefit to RGCs, their effects are often transient and do not prevent the underlying structural instability of mitochondrial membranes that initiates energy loss, oxidative stress, and apoptosis. Therapies that directly stabilize mitochondrial membranes may therefore provide a more robust means of protecting RGCs in glaucoma.

HDAP2 (Biotin-dArg-Phe-Phe-dArg-amide) is a novel high-density aromatic peptide that binds cardiolipin, a mitochondria-specific phospholipid that promotes membrane curvature of the inner mitochondrial membrane. During cellular stress, cardiolipin exhibits phase transition behavior that can destabilize the inner mitochondrial membrane to induce cytochrome c release and mitophagy, a phenomena likely exaggerated in glaucoma ^24^. In contrast to traditional ROS scavengers, HDAP2 is a cardiolipin-specific peptide that binds with high affinity and promotes selective stabilization of the inner mitochondrial bilayer ^25^. This stabilization could potentially prevent membrane rupture and preserve mitochondrial function. In vitro, HDAP2 maintains membrane potentials, suppresses ROS, and promotes cell survival under conditions of high metabolic stress ^25^. In vivo, HDAP2 colocalizes with RGC mitochondria and demonstrated neuroprotective effects in an optic nerve crush mouse model ^26^.

Our hypothesis was that HDAP2 might help maintain mitochondrial integrity in glaucoma and protect RGCs and their axons. To test this, we used HDAP2 to treat DBA/2J mice, a spontaneous glaucoma model in which most eyes develop both high IOP and severe RGC loss. Animals between 4 and 12-months of age were treated every other day with the peptide to test its ability to offer pressure-dependent protection to RGCs and their axons.

## Methods

### Peptide Synthesis

The novel peptide HDAP2 (Biotin-dArg-Phe-Phe-dArg-amide) was synthesized commercially (GenScript, Piscataway, NJ, USA) and purity (>96%) was confirmed by HPLC and mass spectrometry. This peptide targets mitochondria and binds cardiolipin on the inner mitochondrial membrane to maintain the mitochondrial membrane potential during oxidative stress ^25^. HDAP2 is patented by the Research Foundation of the City University of New York.

### Distribution of systemically administered HDAP2 into retina

To assess retinal uptake of HDAP2, adult C57BL/6 mice were injected intraperitoneally with 50 mg/kg biotinylated HDAP2 reconstituted in saline. Animals were sacrificed 2 hours post-injection, and eyes were processed for cryosectioning and immunohistochemistry as described below. Control animals received saline injections only.

Whole eyes were immersed in 4% paraformaldehyde in phosphate buffer (pH 7.3). After 10 minutes, the cornea, lens, and vitreous body were removed to ensure adequate fixative penetration into the retina, and the eyecups were returned to fixative for an additional hour. Following fixation, eyecups were rinsed in buffer and cryoprotected in phosphate-buffered sucrose (10%, 20%, and 30%) prior to embedding in optimal cutting temperature (OCT) medium (Polyfreeze, Polysciences, Warrington, PA). Sections were cut at 14 μm thickness at –25°C using a cryostat (Leica CM3050S) and mounted onto gelatin-coated slides.

HDAP2 uptake was visualized by incubating tissue sections with Alexa Fluor 488-conjugated streptavidin (1:200, Jackson ImmunoResearch, West Grove, PA) overnight at 4°C. Tissues were co-stained with rabbit anti-RBPMS (1:250; PA5-31231, Invitrogen, Carlsbad, CA) to label retinal ganglion cells or rabbit anti-Cox4 (1:200; PA5-29992, Invitrogen, Carlsbad, CA) to label mitochondria. Sections were incubated in secondary antibodies for one hour at room temperature (donkey anti-rabbit conjugated to either Alexa Fluor 488 or 594). ToPro3 (1:1000; Invitrogen, Carlsbad, CA) was used to label cell nuclei. Sections were imaged using an Olympus Fluoview 300 confocal microscope with an Olympus 40× oil immersion objective (NA 1.0). Laser intensity for the 488 nm channel was maintained at ∼530V across all samples to enable direct comparison of streptavidin labeling between treatment conditions.

### Animals and Experimental Design

This was a proof-of-concept study designed to evaluate whether systemic HDAP2 could protect retinal ganglion cells in the DBA/2J mouse model of glaucoma. DBA/2J mice (4 – 8 weeks old, both sexes) were purchased from The Jackson Laboratory (Bar Harbor, ME) and housed at York College, CUNY under standard laboratory conditions. Animals were randomly assigned to control (untreated) or HDAP2-treated groups beginning at 4 months of age. Retinas from control animals were studied at 4 months, before elevated IOP had developed. Mice in the treated group received HDAP2 administered intraperitoneally (IP, 3 mg/kg) every other day from ages 4 through 12 months (∼120 injections per animal). Dose selection was based on tolerance studies, which demonstrated 100% survival at doses below 30 mg/kg with an LD₅₀ of 55 mg/kg ^26^.

Untreated control mice were age– and sex-matched DBA/2J animals from the same colony that were handled only for cage changes and monthly IOP measurements. Because DBA/2J mice are highly sensitive to handling stress and prone to agitation and occasional seizures ^27^, we intentionally did not include a vehicle-injected control group. Repeated handling and saline injections in DBA/2J mice have been shown to induce marked Fos immunoreactivity in neural structures compared with C57BL/6J mice, indicating a heightened strain-specific stress response^28^. Because repeated handling and injections can also transiently elevate IOP in conscious mice ^29^, additional vehicle injections might have introduced variability rather than improved experimental control. Given the scale of the study (100 animals total) and the ethical concern of imposing unnecessary stress, we chose not to subject untreated controls to an eight-month injection regimen. All statistical analyses were adjusted for cumulative IOP exposure, which accounts for longitudinal differences in pressure burden.

IOP was measured in all animals monthly, starting at 4 months of age, using an Icare TonoLab rebound tonometer (Colonial Medical Supply, Franconia, NH) under light isoflurane anesthesia. Since approximately 26% of DBA/2J eyes do not develop high IOP ^30^, if at all, mice studied in the 10-month cohort had eyes that sustained at least 2 months of elevated IOP (>19 mmHg). For all ages studied, we calculated cumulative IOP exposure (cIOP), the sum of monthly IOP values exceeding 19 mmHg, to facilitate correlation between sustained pressure exposure and retinal pathology independent of mean IOP. All procedures were approved by the York College IACUC and conducted in accordance with the ARVO Statement for the Use of Animals in Ophthalmic and Vision Research.

### Tissue Processing and RGC Analysis

HDAP2 treatment effects on RGCs and optic nerves were assessed in 10-, 11-, and 12-month-old animals. Mice were euthanized using an overdose of ketamine (300 mg/kg) and xylazine (60 mg/kg) injected intraperitoneally. Eyes were enucleated, the cornea and lens removed, and the eyecups were immersion-fixed in 4% paraformaldehyde in phosphate buffer for 75 minutes. At enucleation, optic nerves were cut ∼1-2 mm from the globe and fixed in 2.5% glutaraldehyde/2% paraformaldehyde for separate processing.

Fixed retinas were dissected free from the pigment epithelium, rinsed in buffer, and immunostained using standard immunohistochemical procedures ^26^ with rabbit anti-RBPMS (1:250, Invitrogen, cat# PA5-531231, 2-3 days) and Alexa Fluor 488 donkey anti-rabbit (1:500, Invitrogen Cat# A-21206, overnight) antibodies to label RGCs. Retinas were flat-mounted with the ganglion cell layer facing up and coverslipped with Vectashield mounting medium (Vector Laboratories, Burlingame, CA).

Whole retinas were imaged using a Zeiss LSM 900 confocal microscope equipped with Airyscan 3.6 technology and a 10× objective. The imaging plane was focused on RBPMS-labeled somas using 30-50 anchor points and the microscope systematically acquired ∼160 images with 10% overlap. Airyscan processing automatically registered and blended the images to produce a single, high-resolution montage of each retinal wholemount. RGCs counts were obtained using RGCode, a validated deep learning program for automated analysis ^31^, which enabled rapid, unbiased counting of RGCs, with all outputs reviewed by an investigator blinded to the experimental condition.

### Optic Nerve Analysis

Optic nerves were post-fixed in 1% osmium tetroxide for 30 minutes, dehydrated through an ascending series of ethanol, and embedded in a 3:2 mixture of Spurr’s and EPON epoxy resins for 24 hours under vacuum. After polymerization at 60°C for 18 hours, blocks were trimmed and oriented to enable 1 μm thick semithin cross-sections of the nerves cut using a Leica Ultracut ultramicrotome. Sections were mounted on glass slides and stained with toluidine blue on a hot plate for 5 minutes to differentiate myelinated axons from surrounding tissue. Nerves were imaged with a 40× Plan Fluor objective (NA 1.3 oil) using a Nikon Eclipse E800 microscope fitted with an AmScope MU1803 camera.

Individual axons were identified, counted, and measured using the AxonJ plugin for FIJI/ImageJ^32^. The cross-sectional area of each nerve was computed and 4-5 non-overlapping regions (∼0.015 mm² per area) were sampled. Axon counts from the sampled images were extrapolated to predict total axonal content of the nerve. For size comparison, axons were categorized as small (<0.5 μm²), medium (0.5-2.0 μm²), or large (>2.0 μm²) ^33^. To assess nerve quality, each cross-section was graded using a 3-point scale by two investigators blinded to the experimental condition: grade 1 = minimal degeneration (normal honeycomb appearance with well-preserved myelin sheaths); grade 2 = moderate degeneration (partial loss of normal structure with some fiber disorganization); and grade 3 = severe degeneration (extensive axonal loss, cellular debris accumulation, and disrupted tissue organization) ^30,34^.

### Statistical Analysis

Statistical analyses were performed using GraphPad Prism 10 with α = 0.05 for significance testing. Data are presented as mean ± SEM unless otherwise indicated. RGC and axon count data were analyzed using a one-way ANOVA to assess overall treatment effects, followed by Tukey’s multiple comparison post-hoc tests.

We conducted a two-way factorial ANOVA (treatment × pressure level) to evaluate pressure-dependent protective effects. Cumulative IOP values were categorized as low (<40 mmHg), moderate (40-80 mmHg), and high (≥80 mmHg) based on previous studies correlating IOP with retinal damage in DBA/2J mice ^35^. Severe RGC loss was defined as <15,000 RGCs, representing approximately 70% loss, which is consistent with end-stage glaucomatous damage ^36^. Fisher’s exact test assessed associations between treatment condition and severe loss rates. Kaplan-Meier survival analysis determined pressure thresholds for severe RGC loss, with log-rank tests comparing survival curves between treatment groups.

The relationship between cIOP exposure and RGC counts was evaluated using Pearson correlations for each treatment group. Linear regression analysis with slope comparison tests evaluated pressure-response relationships between groups. Axon degeneration scores were analyzed using the Kruskal-Wallis test with post-hoc comparisons. The relationship between RGC and axon counts was assessed using Pearson correlation analysis. Partial eta-squared (η²) was reported for ANOVA effects, and Cohen’s d was calculated for pairwise comparisons to assess biological significance alongside statistical significance.

## Results

### HDAP2 peptide colocalizes with RGCs and Cox4

To determine whether systemically administered HDAP2 reached the retina, we analyzed retinal tissue 2 hours after intraperitoneal injection (see Methods). Endogenous biotin levels in the retina were low in controls (Figure 1A, D). However, administration of biotinylated HDAP2 to DBA/2J mice resulted in peptide detection throughout all layers of the retina, with prominent colocalization in RGCs, indicating that HDAP2 reached RGCs after systemic administration at levels exceeding endogenous biotin (Figure 1B, E). The distribution of HDAP2 overlapped with that of Cox4, suggesting mitochondrial localization (Figure 1C, F).

**Figure 1.**
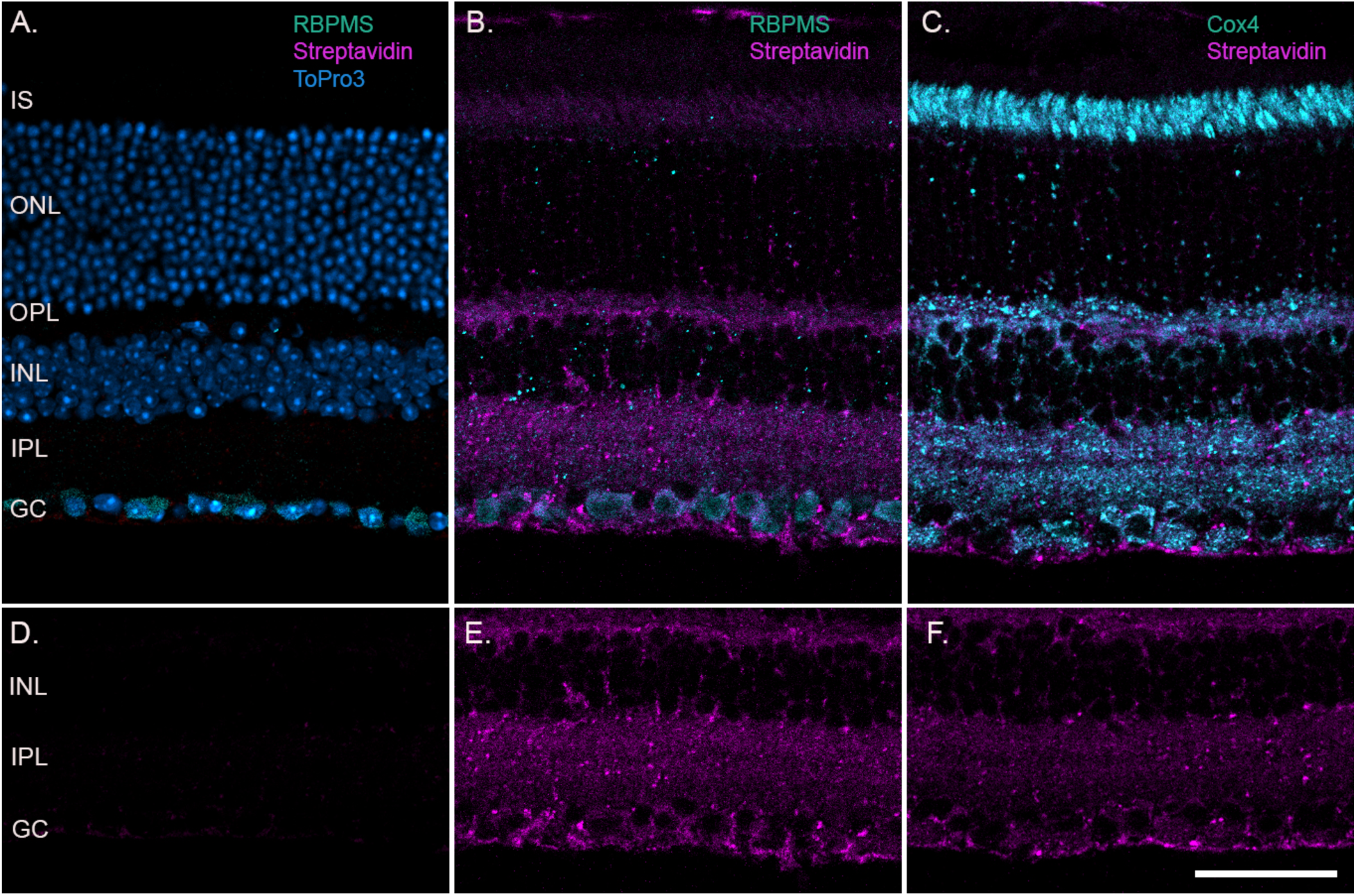
Distribution of systemically administered HDAP2 into retina. Representative retinal cryosections from control and HDAP2-treated animals (50 mg/kg, IP, 2 hours post-injection). **A.** Control retina labeled with streptavidin-conjugated Alexa Fluor 488 (magenta) to detect endogenous biotin, RBPMS (cyan) to identify retinal ganglion cells (RGCs), and ToPro3 (blue) for nuclear labeling. Endogenous biotin levels were very low. **B.** HDAP2-treated retina showing widespread peptide uptake (streptavidin, magenta) with prominent colocalization in RGC somas (RBPMS, cyan) and throughout the plexiform layers. **C.** HDAP2 distribution (streptavidin, magenta) closely matched the mitochondrial marker Cox4 (cyan). **D–F.** Streptavidin-only channel from panels A–C, respectively. Scale bar = 50 µm. GC, ganglion cell layer; PL, inner plexiform layer; INL, inner nuclear layer; OPL, outer plexiform layer; ONL, outer nuclear layer; IS, inner segments of photoreceptors.

### IOP exposure was similar in HDAP2 and untreated groups

We analyzed 62 DBA/2J mouse retinas across three groups: control (n=4), untreated (n=26), and HDAP2-treated (n=32). Control animals were 4 months old and showed no elevated intraocular pressure (IOP). Untreated and HDAP2-treated mice were 10-12 months old with varying IOP exposures. IOP continued to rise over time in both groups (Figure 2A), though individual pressures were highly variable. Total IOP burden, measured by area under the curve (AUC), was not statistically different between groups (Untreated: 139.6 ± 11.96 versus HDAP2: 137.9 ± 11.11 AUC units; p = 0.84, unpaired t-test, Figure 2B), confirming that both experimental groups were exposed to a similar range of ocular pressures.

**Figure 2.**
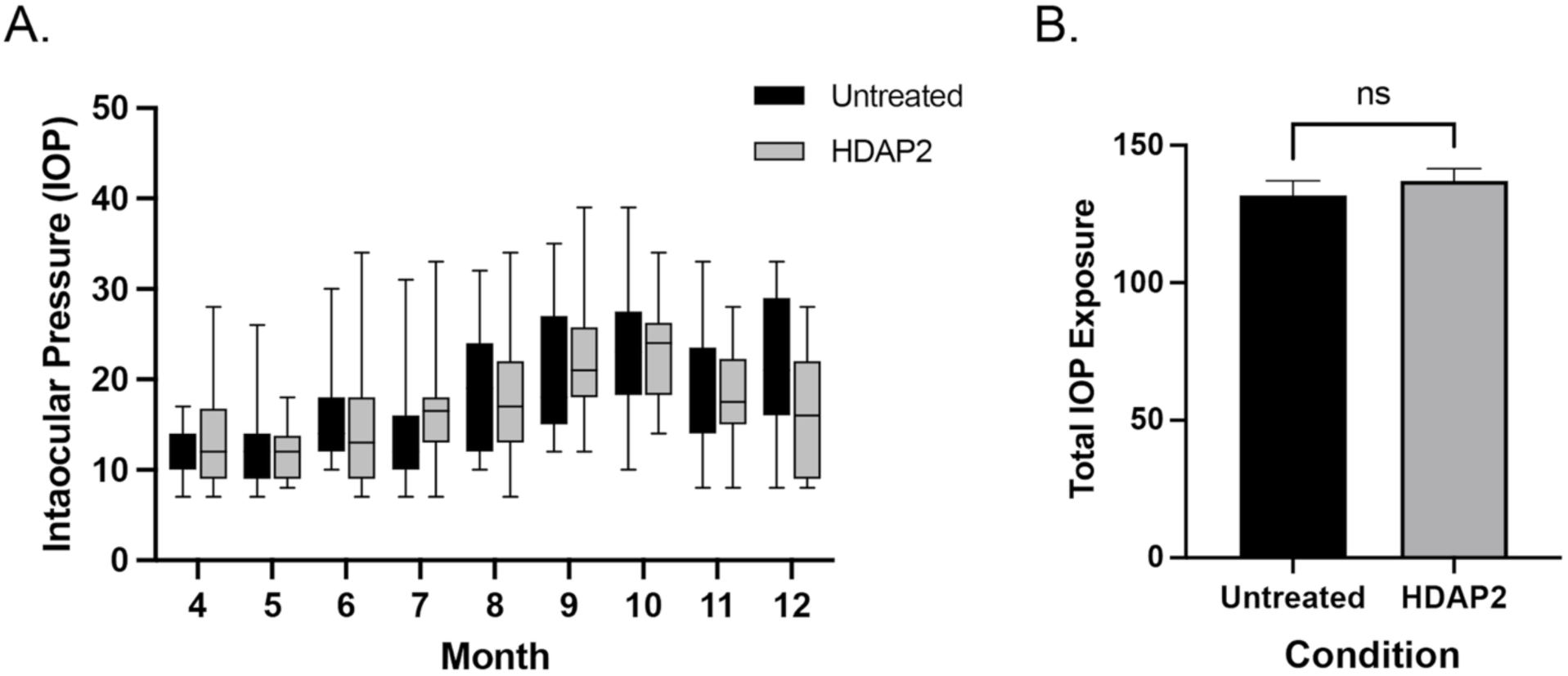
IOP in HDAP2– and control-treated DBA/2J mice. (A) IOPs measured monthly from 4 to 12 months of age. Pressures increased over time in both groups, with high inter-individual variability. (B) Total IOP exposure, defined as the area under the curve representing IOP over time, was not significantly different between groups (p = 0.84, unpaired t-test).

### HDAP2 treatment preserved RGC survival

Qualitative differences in RGC labeling were evident across untreated, control and HDAP2 groups (Figures 3 and 4). Young control retinas that were not exposed to elevated IOP, showed the most uniform RBPMS labeling, with the highest densities in the center and lowest in periphery. This pattern was observed in nearly all retinas from untreated and HDAP2 groups at any age (Figures 3B and C, left; Figure 4A and B, left). The most dramatic differences in RBPMS labeling occurred among untreated animals at moderate and high-pressure exposures (Figures 3B and 4A). In these retinas, many were completely devoid of RBPMS labeling or contained triangular sectors of spared cells. In contrast, retinas from HDAP2 treated animals appeared similar to controls at moderate pressures and only showed damage at high pressures in the 12-month group, where sectors of missing RBPMS label were occasionally observed (Figures 3C and 4B).

**Figure 3.**
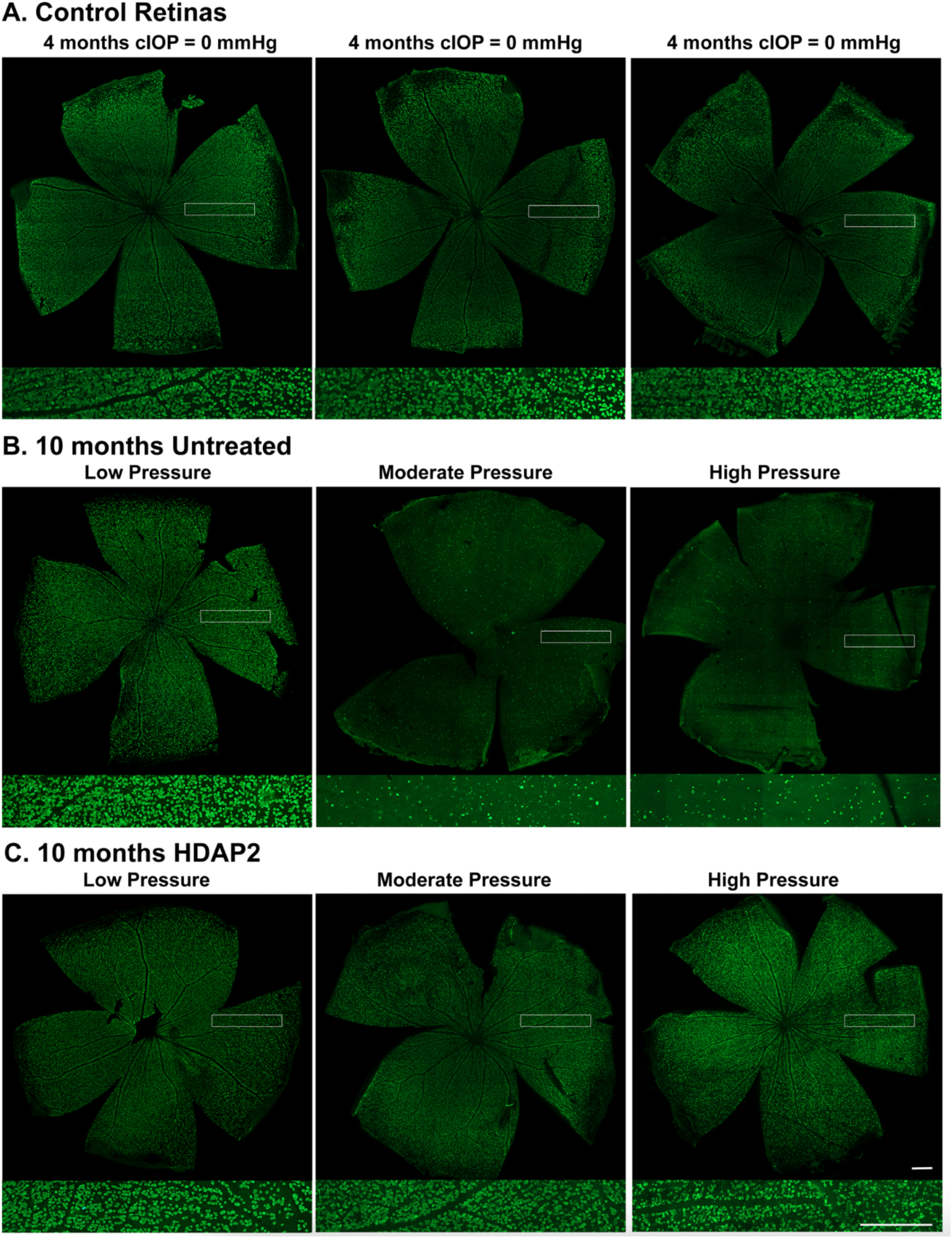
Retinal wholemounts from 4-month-old controls and 10-month-old animals. Wholemounts showing RBPMS-labeled RGCs from control animals and retinas from untreated and HDAP2-treated eyes at different cIOPs. Pressure categories: Low (cIOP < 40 mmHg), Moderate (cIOP 40–80 mmHg), and High (cIOP > 80 mmHg). Higher magnification insets show representative RBPMS labeling. (A) Retinas from 4-month-old animals (cIOP = 0 mmHg) showed normal RGC densities. (B) Untreated retinas showed normal RGC distributions at low pressure but progressive RGC loss with moderate and high IOP exposure. (C) HDAP2-treated retinas retained near-normal RGC density even at high pressure (cIOP = 116 mmHg). Scale bars = 500 µm.

**Figure 4.**
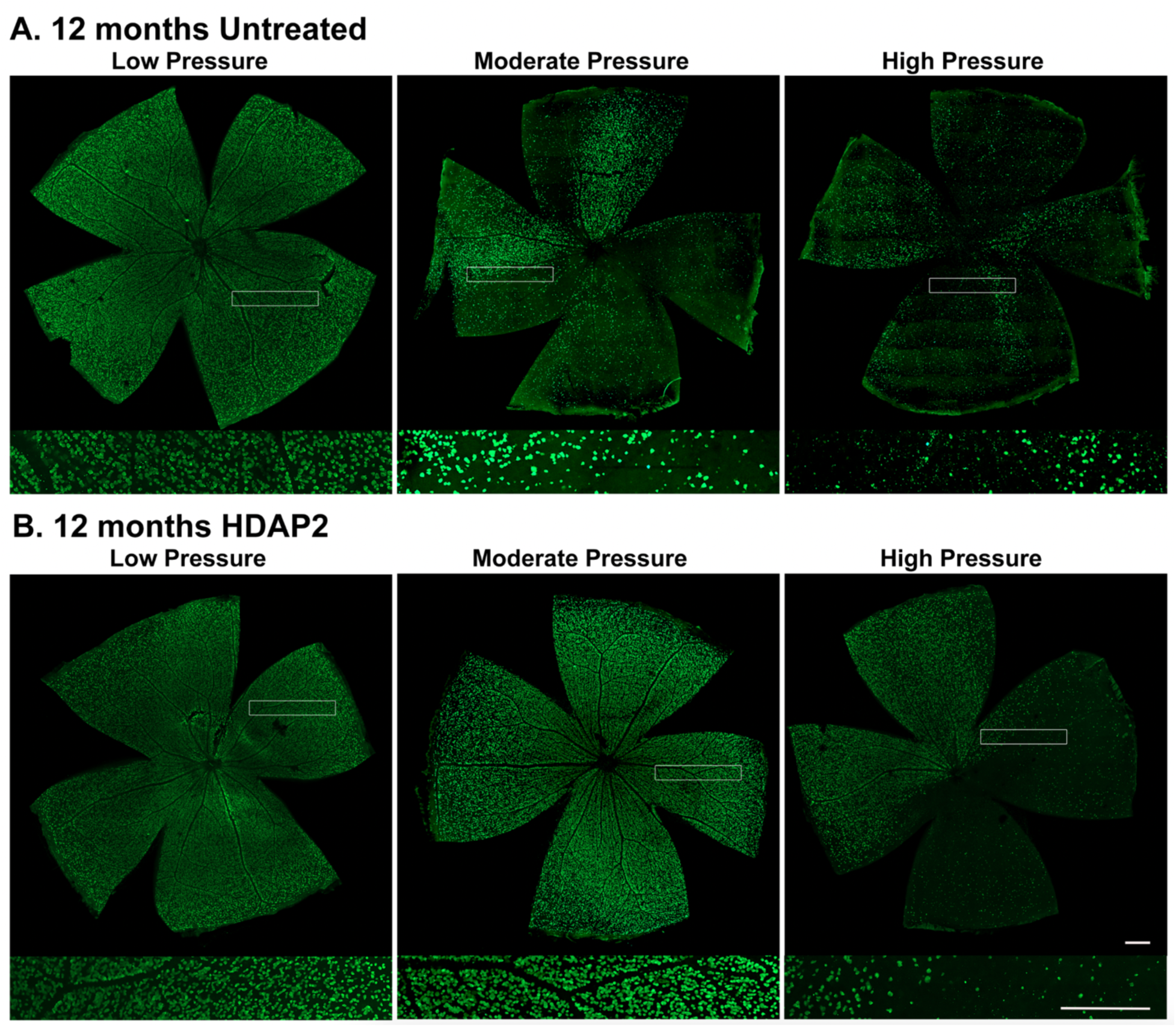
Retinal wholemounts from 12-month-old animals. RBPMS-stained wholemounts to show distribution of RGCs in untreated and HDAP2-treated eyes; higher magnification insets show representative labeling. Pressure groups: Low (cIOP < 40 mmHg), Moderate (cIOP 40–80 mmHg), and High (cIOP > 80 mmHg). (A) Untreated retinas exhibited prominent RGC loss at moderate (cIOP = 75 mmHg) and high pressures (cIOP = 85 mmHg). (B) Retinas from 12-month-old HDAP2-treated eyes maintained higher RGC densities compared to untreated eyes at similar pressures (moderate cIOP = 52 mmHg; high cIOP = 103 mmHg). Scale bars = 500 μm.

RGC counts in the three groups supported these qualitative observations. Retinal wholemounts showed that HDAP2-treated retinas across all age groups and pressure exposures had similar numbers of RGCs compared to control retinas (Table 1, Figures 3 and 4). Control retinas averaged 51,727 ± 5,963 RGCs, untreated retinas averaged 26,419 ± 4,017 RGCs, and HDAP2-treated retinas averaged 38,842 ± 2,789 RGCs. A one-way ANOVA revealed a highly significant overall treatment effect on RGC survival (F(2,59) = 5.524, p = 0.0063, R² = 0.158). Post-hoc comparisons demonstrated that untreated retinas lost 25,308 RGCs relative to controls (p = 0.012, Cohen’s d = 1.42), while HDAP2-treated retinas preserved 12,423 more RGCs than untreated eyes (p = 0.015, Cohen’s d = 0.70). HDAP2-treated retinas did not differ significantly from controls (p = 0.12, Cohen’s d = 0.73). Overall, HDAP2 treatment preserved approximately 49% of RGCs that were otherwise lost in untreated controls under equivalent pressure exposures.

**Table 1:** RGC Survival.

| Comparison | Mean Difference<br>(RGCs) | 95% CI | p-<br>value | Cohen's<br>d | Effect<br>Size |
| --- | --- | --- | --- | --- | --- |
| Control vs<br>Untreated | +25,308 | [7,843 to 42,772] | 0.012* | 1.42 | Very large |
| Untreated vs<br>HDAP2 | -12,423 | [-22,264 to -<br>2,582] | 0.015* | 0.70 | Medium |
| Control vs HDAP2 | +12,885 | [-4,707 to<br>30,477] | 0.12 | 0.73 | Medium |

\* $p < 0.05$ ; Overall one-way ANOVA: $F(2,59) = 5.524$ , $p = 0.0063$ , $R^2 = 0.158$ .

### HDAP2 protection was most apparent at high IOP

Given the large variability in IOP measurements across animals, we compared cIOP to surviving RGCs using linear regression analysis (Figure 5). Control animals with high RGC counts are included for reference. Untreated retinas showed strong, pressure-dependent RGC loss (slope = – 404.7 RGCs/mmHg, R² = 0.42, p = 0.0004), while HDAP2-treated retinas lost cells more slowly with less than half the pressure sensitivity (slope = –254.9 RGCs/mmHg, R² = 0.28, p = 0.0018). However, the difference between the slopes was not statistically significant (F(1,54) = 1.54, p = 0.22).

**Figure 5.**
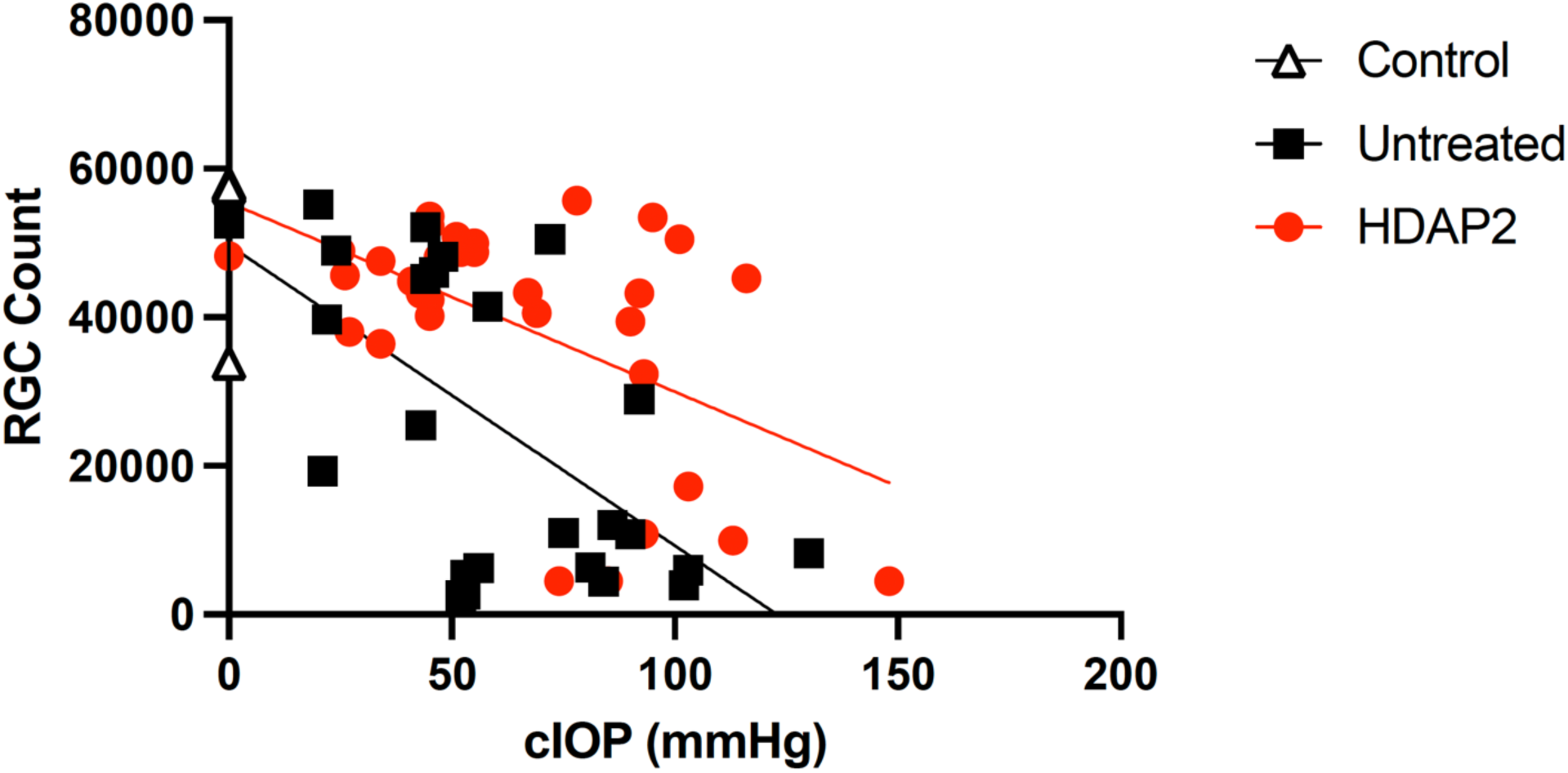
Linear regression of RGC survival versus cIOP. Regression lines were fitted for untreated and HDAP2-treated eyes versus IOP, with control eyes plotted for reference. Untreated eyes demonstrated rapid RGC loss with increasing IOP (slope = –404.7, 95% CI –606 to –203; R² = 0.42, p = 0.0004). HDAP2-treated eyes also showed decline but significantly more slowly (slope = –254.9, 95% CI –407 to –103; R² = 0.28, p = 0.0018). The extrapolated x-intercepts suggested that HDAP2-treated eyes endured pressure over an extended range (217.6 mmHg) compared to untreated eyes (122.9 mmHg). There were no significant differences between the slopes (F(1,54) = 1.54, p = 0.22).

To further examine pressure effects, we grouped animals by low (<40 mmHg), moderate (40-80 mmHg), and high (>80 mmHg) cIOP exposure levels (Figure 6). A two-way ANOVA revealed a significant interaction (F(6,55) = 6.99, p < 0.0001), suggesting that HDAP2 efficacy depends on IOP severity. Under mild pressure exposure, both experimental groups showed minimal RGC loss (44,146 ± 2,963 HDAP2 vs. 44,914 ± 5,505 untreated), without significant deviation from 4-month-old controls (51,727 ± 5,963). Under moderate pressure, untreated animals suffered substantial RGC loss while HDAP2-treated animals maintained significantly better survival (44,453 ± 3,109 vs. 28,057 ± 6,070; 58% improvement). At high pressure exposures, HDAP2-treated animals retained 28,298 ± 5,776 RGCs versus 10,092 ± 2,876 in untreated eyes (180% improvement). Tukey’s post-hoc multiple comparisons tests confirmed significant protection at moderate and high pressures (p < 0.05).

**Figure 6.**
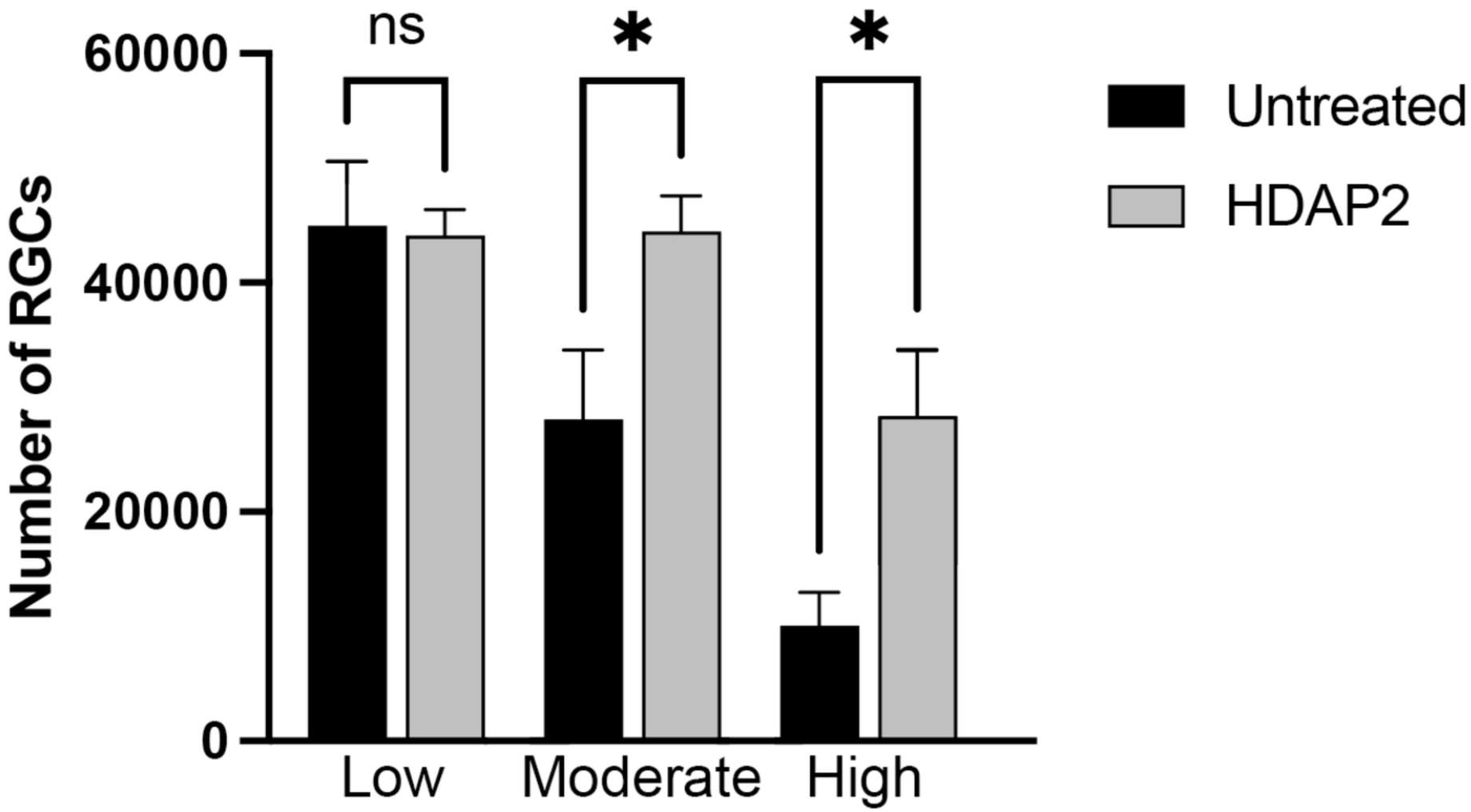
RGC survival according to pressure exposure in untreated and HDAP2-treated eyes. Bar graph shows mean RGC counts (±SEM) at low (<40 mmHg), moderate (40–80 mmHg), and high (≥80 mmHg) cIOP. Black bars: untreated eyes; gray bars: HDAP2-treated eyes. Two-way ANOVA revealed a significant treatment × pressure interaction (F(6,55) = 6.99, p < 0.0001) but no main effects. Post-hoc Tukey tests revealed no significant differences between treatments at low pressure, whereas HDAP2 resulted in significantly higher numbers of RGCs under moderate and high pressures compared to untreated eyes. ns = non-significant; *p < 0.05.

The pressure-dependent differences in RGC survival redefined the basic IOP thresholds for RGC pressure tolerance across the treatment groups. We defined severe RGC loss as <15,000 RGCs (approximately 67-70% loss compared to age-matched controls), a level roughly equivalent to severe glaucomatous damage ^36,37^. Kaplan-Meier survival analysis indicated that the median cIOP for reaching this threshold was 84 mmHg (95% CI 60-107 mmHg) in untreated eyes and 113 mmHg (95% CI 75-162 mmHg) in HDAP2-treated eyes (Figure 7A), which represents a 29 mmHg increase in pressure tolerance (Log-rank test: χ² = 8.724, p = 0.003). Consistent with this finding, Fisher’s exact test revealed that untreated groups were more susceptible to severe RGC loss (46.2%, 12/26) than HDAP2-treated animals (15.6%, 5/32; p = 0.019, OR = 4.63), which represents a clinically significant reduction in risk (Figure 7B).

**Figure 7.**
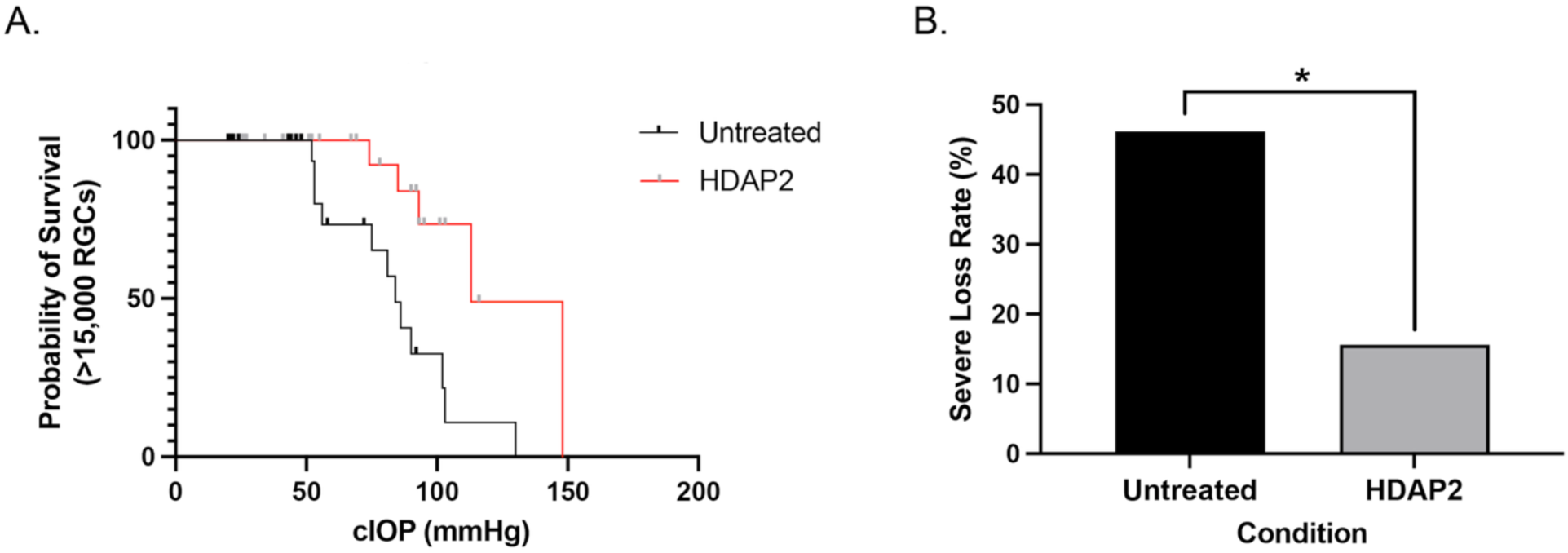
Pressure thresholds for severe RGC loss in untreated and HDAP2-treated eyes. (A) Kaplan-Meier survival curves showing eyes retaining >15,000 RGCs across cIOP exposure. HDAP2-treated eyes (red) tolerated 29 mmHg of additional pressure before showing severe loss compared to untreated eyes (black) (Log-rank χ² = 8.724, p = 0.003). (B) Severe loss rates (<15,000 RGCs): Untreated 46.2% (12/26) vs. HDAP2 15.6% (5/32). Fisher’s exact test: p = 0.019, OR = 4.63. *p < 0.05.

### HDAP2 preserved optic nerve axons

To study whether HDAP2 also protected optic nerves, we analyzed cross-sections from 4-month-old controls and 10-month-old mice with similar pressure exposures (control: n=6; untreated: n=6; HDAP2: n=5) (Figure 8). Semithin cross-sections of HDAP2-treated nerves showed remarkably preserved morphology that resembled the normal nerve structure from controls. This contrasted significantly with the severe degeneration observed in untreated optic nerves, which presented with massive axonal degradation, disrupted myelin, and compromised architecture (Figure 8A).

**Figure 8.**
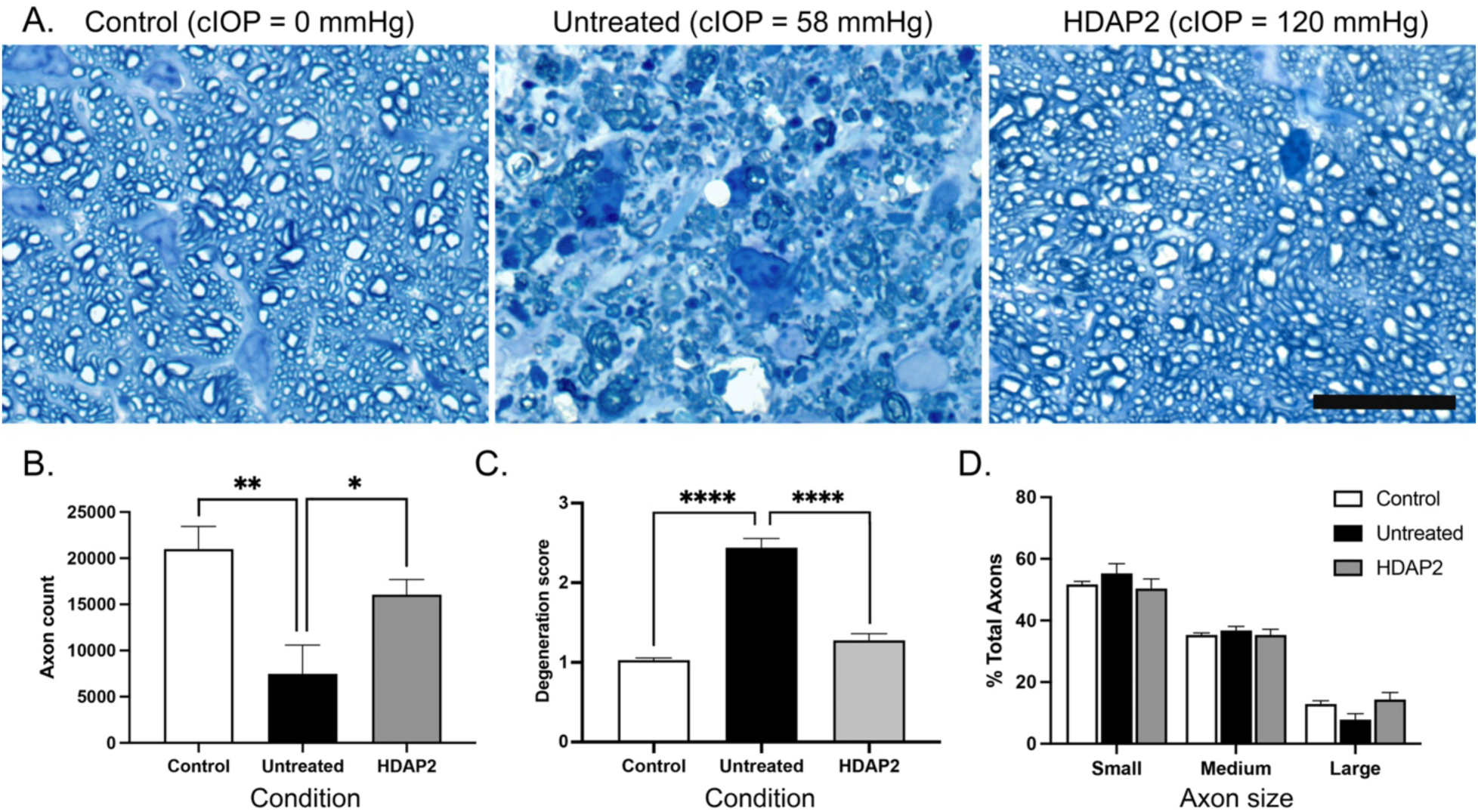
Optic nerve morphology, axon numbers, quality scores, and size distributions. (A) Semithin cross-sections from 4-month-old controls and 10-month-old experimental animals. Untreated nerves showed extensive axonal degeneration and myelin disruption, while HDAP2-treated nerves maintained preserved honeycomb-like architecture similar to controls. Scale bar = 20 μm. (B) Axon counts per cross-section. Control nerves: 20,984 ± 2,464 axons; HDAP2-treated: 16,047 ± 1,657; untreated: 7,476 ± 3,109. One-way ANOVA: F(2,14) = 7.426, p = 0.0063, R² = 0.515. (C) Axon quality scores showed significant group differences (Kruskal-Wallis χ²(2) = 62.86, p < 0.0001, ε² ≈ 0.59). Dunn’s post-hoc tests revealed that untreated nerves were significantly more degenerated than controls (p < 0.0001) and HDAP2-treated nerves (p < 0.0001), while HDAP2-treated and control nerves did not differ (p = 0.4010). (D) Axon size distribution showing mean ± SEM proportions of small (<0.5 μm²), medium (0.5–2.0 μm²), and large (>2.0 μm²) axons. Two-way ANOVA: size effect F(2,42) = 331, p < 0.0001; no treatment effect F(2,42) = 0.00, p = 0.9998; no interaction F(4,42) = 2.5, p = 0.0545. HDAP2 maintained normal axon size distribution, suggesting unbiased protection across RGC subtypes.

Extrapolated counts revealed that control nerves averaged 20,984 ± 2,464 axons, HDAP2-treated nerves had 16,047 ± 1,657 axons, and untreated nerves had only 7,476 ± 3,109 axons per cross-section (Figure 8B). A one-way ANOVA confirmed highly significant group differences (F(2,14) = 7.426, p = 0.0063, R² = 0.515). Post-hoc tests indicated that untreated nerves had significantly fewer axons than both controls (p = 0.0072) and HDAP2-treated nerves (p = 0.0430), while control and HDAP2-treated groups did not differ significantly (p = 0.1332).

Qualitative comparison showed that HDAP2 maintained axon morphological quality of the nerves compared to untreated controls (Figure 8C). There was a significant main effect of group on degeneration scores (Kruskal-Wallis χ²(2) = 62.86, p < 0.0001, ε² ≈ 0.59). Dunn’s post-hoc tests revealed that degeneration in untreated nerves differed highly significantly from both controls (p < 0.0001) and HDAP2-treated nerves (p < 0.0001), whereas controls were not statistically different from HDAP2-treated nerves (p = 0.4010). Axon preservation correlated positively and significantly with RGC survival across animals (r = 0.65, p = 0.016), suggesting that HDAP2 protected both cell bodies and their axonal projections.

### HDAP2 preserved axon size distribution across RGC subtypes

To assess whether HDAP2 provided comprehensive RGC protection versus preferential protection of specific subsets, we analyzed axon size distribution as a measure of RGC functional diversity (Figure 8D). Axons were categorized as small (<0.5 μm²), medium (0.5-2.0 μm²), or large (>2.0 μm²) based on cross-sectional area, to represent structural correlates of different RGC functional classes ^33^. A two-way ANOVA (treatment × axon size) revealed a robust main effect of axon size (F(2,42) = 331, p < 0.0001), as expected given the natural bias toward small axons, but no treatment effect (F(2,42) = 0.00, p = 0.9998) and no interaction (F(4,42) = 2.5, p = 0.0545). This conservation of normal size distribution suggests that HDAP2 confers nonselective neuroprotection to all RGC types rather than selectively preserving specific RGC subtypes.

## Discussion

This study demonstrates that HDAP2, a cardiolipin-targeted peptide, provided robust neuroprotection in the DBA/2J model of glaucoma. HDAP2 preserved both RGCs and optic nerve axons despite sustained exposure to elevated IOP, with the strongest relative protection observed under the highest pressures. Treated retinas retained nearly half of the RGCs that were otherwise lost in untreated animals, and optic nerve morphology in HDAP2-treated eyes resembled those of controls. These results provide proof-of-concept that stabilizing cardiolipin to maintain mitochondrial membrane integrity can substantially increase RGC tolerance to elevated IOP in glaucoma.

### HDAP2’s protection of RGCs became most apparent at elevated IOP

RGC survival at low IOPs was comparable to control retinas in both experimental conditions. At moderate and high IOPs, however, HDAP2 showed clear protection, with RGC survival 58% greater at moderate exposure and 180% at high pressures. These results are consistent with HDAP2’s mechanism. The peptide binds with cardiolipin to stabilize mitochondrial membranes. Membrane stabilization should occur at all conditions, although HDAP2’s benefit will only be apparent when cellular stress triggers events that lead to mitochondrial dysfunction (i.e., cardiolipin phase transitions, cytochrome c release, and mitochondrial depolarization). Although mitochondrial potential, ROS, or cardiolipin binding were not directly measured, our companion work has shown that HDAP2 binds cardiolipin, stabilizes its bilayer phase, and inhibits oxidative stress in cultured cells ^25^ and supports RGC survival following optic nerve crush ^26^. Together, these data support the interpretation that HDAP2 preserves RGC survival by maintaining cardiolipin integrity and mitochondrial function.

### Comparison with other mitochondrially-directed compounds

Our results suggest that HDAP2 acts through a mechanism fundamentally distinct from other mitochondria-targeted therapies. Agents such as SS-31 ^21^, SK-Q ^38^, and MitoQ ^39^ primarily act as antioxidants, while nicotinamide supports NAD⁺ metabolism ^20,40^. Although these compounds offer partial protection of RGCs in animal models, their benefits are typically limited, both transient in duration and inconsistent across disease stages. They also act downstream of the initial mitochondrial instability, and do not prevent the early events that trigger degeneration. Mitochondrial transplantation has also been explored but faces significant practical challenges ^41^. In contrast, HDAP2 acts upstream by binding directly to cardiolipin to stabilize the inner mitochondrial membrane, thereby intervening before apoptotic signaling cascades are triggered. This action likely underlies the resilience observed, even under cIOP elevations exceeding 100 mmHg. By shifting the threshold for severe RGC loss upward by nearly 30 mmHg, HDAP2 produced a neuroprotective effect that is comparable in magnitude to conventional IOP-lowering therapies, but achieved through direct neuroprotection rather than pressure reduction.

### Clinical implications

These results have important clinical implications. Current glaucoma therapies focus almost exclusively on lowering IOP, yet many patients continue to lose vision even when pressure is adequately controlled. Standard treatments typically achieve only a 20–30% reduction in IOP, corresponding to about 4–8 mmHg ^42–44^. By contrast, HDAP2 shifted the threshold for severe retinal ganglion cell (RGC) loss upward by 29 mmHg, a level of protection comparable in magnitude to traditional therapies but achieved through direct neuroprotection rather than pressure reduction. This effect translated into a 4.6-fold reduction in the risk of severe visual impairment (46.2% vs. 15.6%), a difference with clear clinical relevance.

The ability of HDAP2 to preserve RGCs at elevated IOP highlights cardiolipin stabilization as a potential adjunct to pressure-lowering approaches. Such a strategy may be particularly beneficial for patients with normal-tension glaucoma or treatment resistant disease, where reducing IOP alone is insufficient to prevent progression. In this study, we found that treated eyes tolerated pressure levels that would otherwise produce extensive RGC loss. Importantly, HDAP2 protected both RGC somas and axons, and its benefit extended across RGC subtypes, which raises the prospect of broad preservation of visual function.

### Limitations and future directions

While our findings show compelling evidence for HDAP2’s efficacy in protecting RGCs and axons from elevated IOP, we recognize that more work is needed. For this proof-of-concept study, the peptide was administered intraperitoneally to control for dosing. Because DBA/2J mice are unusually sensitive to handling stress and prone to agitation and seizures, we did not include a vehicle-injected control group. Chronic injections over eight months would have imposed unnecessary stress on untreated animals. Although handling-related stress is not expected to confer neuroprotection, we acknowledge this as a study limitation, and future confirmatory work will include vehicle-treated controls.

For therapeutic application, alternative delivery routes, such as topical, nasal, intravitreal, or sustained-release formulations, will need to be developed. In addition, our current data set did not allow us to distinguish the relative effects of acute high-pressure spikes, chronic pressure elevation, and age-related vulnerability, which will be helpful for understanding limitations of this treatment. It appeared that high-pressure spikes and age were greater risk factors for RGC death, but our sample sizes were not large enough for us to draw conclusions.

Finally, while the results strongly suggest a mitochondrial mechanism for HDAP2’s action, direct evidence of cardiolipin stabilization in DBA/2J mice is still needed. Ongoing work using electron microscopy to evaluate mitochondrial morphology, as well as functional testing, will help address these questions.

## Acknowledgments

We thank the Glaucoma Research Foundation (ShafGrnt2024MacNM), and NIGMS R16GM149428 (M.A.M.) for funding support, Ms. Karen Manifold and Ms. Sara Arain for excellent animal care, and the Biology Department at Queens College, CUNY for use of their confocal microscope

## Funding

This work was supported by grants from the Glaucoma Research Foundation (ShafGrnt2024MacNM) and NIGMS R16GM149428 (MAM).

## Commercial Relationships Disclosures

MAM: Inventor on a patent related to HDAP2 therapeutic applications (Research Foundation, CUNY); no current commercial relationships; WM: None; AB: Inventor on a patent related to HDAP2 therapeutic applications (Research Foundation, CUNY); no current commercial relationships.

## Notes

### Competing Interest Statement

The authors have declared no competing interest.

